# Default(y) Mode Network: Important regions of DMN do not survive alterations in flip angles

**DOI:** 10.1101/2020.07.09.196568

**Authors:** Ishan Singhal, Abhishek K. Soni, Narayanan Srinivasan

## Abstract

The default mode network (DMN) is thought to capture intrinsic activity of the brain and has been instrumental in understanding the dynamics of the brain. However, the DMN has not been without critics; both conceptual and empirical. The empirical criticisms caution against physiological noise as a source for the reported connectivity in the DMN. Smaller flip angles (FAs) have been modelled and shown to reduce physiological noise in BOLD signal recordings. A previous functional MRI (fMRI) study with flickering checkerboard stimuli, manipulated FAs to show that activity in the posterior-cingulate cortex (PCC) and precuneus is prone to physiological noise. This raises questions about studies that show activations in these areas (PCC and precuneus) with a fixed FA and the role of these areas in brain networks like DMN. Given the prominent role of PCC and precuneus in DMN, we studied the effect of FAs on the resting-state functional connectivity involving these areas in DMN. We used four FAs and recorded resting-state activity in a 3-T scanner. The results show PCC and precuneus BOLD functional connectivity is inconsistent. We lend support to previous empirical criticisms of DMN, linking its activity to physiological noise. Our results add to concerns about PCC and precuneus related BOLD activity and their putative role in DMN. Alongside previous studies we advocate using smaller flip angles as an empirical tool to investigate physiological noise in fMRI studies.

## Introduction

The formalization of a default mode network has been a useful tool in understanding the underlying neural activity when a person is not performing an explicit task (Raichle et al., 2001, Greicius et al., 2003). It has been claimed to capture the intrinsic neural activity of the brain (Raichle & Mintun, 2006). This activity has been used as a baseline measure of neural activity in comparison to tasks (Gusnard & Raichle, 2001) and to monitor how this particular network behaves dynamically when switching from rest to task (Greicius, Krasnow, Reiss & Menon, 2003; Fox et al., 2006). Moreover, deviations in this intrinsic activity has been shown to be useful in diagnosing various clinical conditions (Anticevic et al., 2012).

However, the notion of DMN has not been without controversy and has drawn both conceptual and empirical criticisms. Conceptually, the DMN faces scepticism towards what it means to be at rest, and whether there is anything like a “resting state” or a “default mode” at all and additionally whether it can be considered a true baseline (Morcom & Fletcher, 2007a; 2007b).

Its empirical and instrumental successes are also not without critics. It has been argued that the associated activity in the DMN is confounded with physiological artefacts (Shmueli et al., 2007; Munck et al., 2008; Murphy, Birn & Bandettini, 2013). For instance, activations in the regions near posterior cingulate cortex (PCC) have been linked to noise confounds from both cardiac and respiratory artefacts as well as CSF fluctuations (Wise et al., 2004; Birn et al., 2006; Bianciardi et al., 2011). These concerns perhaps hold stronger ground when studies recording EEG and BOLD signals simultaneously, fail to find correspondence for the resting state network regions (Yuan et al., 2012).

All fMRI studies have to deal with the problem of physiological noise confounds. The problem is more severe with the use of stronger magnetic fields in the scanner since the interaction between physiological noise and the BOLD signal is multiplicative (Wald & Polimeni, 2017). One possible solution for this includes recording cardiac and breathing cycles and using them as nuisance regressors when modelling BOLD time series (Birn et al., 2009). Another suggested solution has been to use a variety of flip angles lower than the suggested Ernst angle (Gonzalez-Castillo et al., 2011; Renvall, Nangini & Hari, 2014; Wald & Polimeni, 2017).

### Current Study

A standard amongst cognitive neuroscientists and neuroscientists alike has been to set the flip angle (FA) above 77° (between 80 and 90), while collecting BOLD data. This value has been prescribed and commonly used since due to arguments that the signal-to-noise ratio is better at angles above 77° (Gonzalez-Castillo et al., 2011). However, this is true only when the major source of noise is thermal but not physiological. When, physiological noise is the major contributor to total noise, lower flip angles provide similar signal-to-noise ratios and are less sensitive to physiological artefacts (Kruger & Glover, 2001; Gonzalez-Castillo et al., 2011). In addition, they also allow lower deposition of heat by reducing radio-frequency power, higher tissue contrast, and lesser head motion related artefacts. This makes smaller FAs an excellent tool for empirically investigating the contributions of cardiac, respiratory and motion related artefacts to BOLD signals (Gonzalez-Castillo et al., 2011).

A previous study (Renvall, Nangini & Hari, 2014) did exactly this; they showed participants a flickering checkerboard inside a scanner and recorded BOLD activity under various flip angle conditions. Their results show that BOLD activity in certain brain regions reduces and eventually disappears as the flip angles are lowered, while other regions remain unaffected. For instance, they report that when FAs are lowered, BOLD activity in PCC, precuneus and posterior superior temporal gyrus (pSTG) regions of the brain are extremely unreliable (i.e. inconsistent across FAs and their activity decreasing/disappearing at some FAs). However, BOLD activity in regions like primary visual cortex V1, lingual gyrus and middle occipital gyrus remains unaffected by changes in FAs. They explain their results by attributing the recorded activity in PCC, precuneus and pSTG to physiological noise, since they show activation only at higher FAs. The authors are wary about the implications of their results on literature in resting-state and DMNs and they write “*PCC and precuneus are nodes of the ‘‘default-mode’’ resting-state network … In our data, PCC and precuneus showed prominent non-BOLD effects, meaning that the default-mode signal decrease in fMRI might include non-BOLD effects*.” (Renvall, Nangini, & Hari, 2014, p. 5).

We conducted our study to directly examine this claim. It has been shown theoretically and experimentally that low FAs can reduce the degree of physiological noise in BOLD signals under certain conditions (Gonzalez-Castillo et al., 2011). Given the empirical criticisms against DMN arguing that its activity is affected by physiological artefacts, we investigated the claims about resting state DMN under different FA conditions.

We recorded resting state activity in healthy, adult participants under different FA conditions to examine whether the DMN activity is prone to alterations in FAs. If the argument that physiological noise influences PCC and precuneus, which are considered the hubs of DMN and changes in flip angles alter the influence of physiological noise are true, then changes in flip angles would lead to smaller functional connectivity indices in PCC and precuneus, but not in medial frontal and prefrontal regions (mPFC) or lateral parietal (LP) regions. This prediction is directly based on results from an earlier task-based study manipulating FAs (Renvall et al., 2014), but also from other resting-state studies linking activity in the DMN to cardiac, respiratory, CSF-related and vascular artefacts (Shmueli et al., 2007; Munck et al., 2008; Murphy, Birn, & Bandettini, 2013). The PCC and precuneus are considered critical components of DMN to the extent that they are most frequently used to identify the DMN in an informed seed-based analysis (Fransson & Marrelec, 2008; Utevsky, Smith & Huettel, 2014). Therefore, we predicted a reduction in functional connectivity indices in PCC and precuneus with decrease in FAs due to reduction in physiological noise artefacts in the BOLD signal.

## Method

### Participants

Nineteen participants from the University of Allahabad campus volunteered and participated in the study. Two participants did not complete the study. Data from one participant was excluded from analysis due to excessive head motion movement. Data from the remaining 16 participants (mean age = 22.81, 6 females) was analysed. Participants were healthy adults and did not self-report any history of serious medical or psychiatric illnesses. Participants took part in the study after granting informed consent and were debriefed after the experiment. The study was approved by the Institutional Ethics Review Board of the University.

### Procedure

Each participant underwent four consecutive eyes-closed resting state fMRI scan sessions. Each session lasted for four minutes and had a different flip angle (FA15°, FA50°, FA77° and FA90°). To rule out order effects, each participant completed the experiment in a unique order of FAs. In total, participants spent approximately 25 minutes inside the scanner. The initial 5-7 minutes were spent in collecting structural scans and gradient echo (GRE) field mapping and the rest of the time was spent undergoing consecutive resting state BOLD activity functional scans. Participants were given an emergency buzzer to use in case they felt any discomfort and wanted to abort the experiment.

Participants were asked to lay inside the scanner with their eyes closed. Participants were asked to remain as still as possible during the scanning. They were asked not to ruminate on any particular topic or thought and were told not to do repetitive tasks like counting. They were also asked to try and avoid falling asleep. At the end of each block, participants were told how many sessions were remaining. This was also informally used as a test by the experimenter to ensure participants had not fallen asleep.

### Data Acquisition

The scans were acquired on a long-bore 3-T Skyra, Siemens whole-body scanner (Version: Syngo MR E11, Siemens, Germany) and a standard 32 channels head coil. Structural MRI images were acquired through two T1-weighted magnetic resonance (MR) images, magnetization-prepared rapid-acquisition gradient echo (MPRAGE; TR = 2000 ms, TE = 2.29 ms, FA = 8°, slice per slab = 176, FoV = 240 mm, Matrix 256×256 TI = 900ms, voxel size = 0.9×0.9×1.0 mm^3^). Resting-state functional images were acquired using an echoplanar (EPI) sequence (TR = 2500 ms, echo time TE = 30 ms, flip angle = 15°, 50°, 77° and 90°, field of view (FoV) 192 mm, matrix = 96 × 96, slice thickness = 3.0 mm, voxel size = 2.0×2.0×3.0 mm^3^ for measurement of BOLD signal). In order to reduce noise from the scanner, noise attenuating headphones were used. To restrict head motion, foam pads on both sides of the head were placed inside the coil.

### Pre-processing

The data pre-processing steps were performed using the functional connectivity toolbox CONN (https://www.nitrc.org/projects/conn) (Whitefield-Gabrieli & Nieto-Castanon, 2012). Statistical Parametric Mapping (SPM 12, Wellcome Department of Imaging Neuroscience, University College London, U.K.) was used for brain imaging data in a separate/verify pre-processing analysis. We used the CONN toolbox since it was specifically designed for functional connectivity analysis and uses a computation-based noise correction strategy (CompCor method) to deal with physiological noise specifically with the resting state network. As such CONN was an ideal analysis platform for our purposes. Standard pre-processing pipelines for volume-based analysis was performed for each individual subject. Functional volumes for each participants’ data underwent realignment and unwarp (head motion estimation and correction) followed by slice-timing correction and normalization. The structural images and functional volumes were normalized to MNI (Montreal Neurological Institute) space using the EPI (echo-planar imaging) template. The images were smoothed using a Gaussian kernel of 8 mm at Full Width at Half Maximum. In order to segment anatomical and functional images into cerebrospinal fluid (CSF) and white matter (WM), subjectlJspecific CSF and WM templates were generated using the SPM12 unified segmentation and normalisation procedure. These images were co-registered for each subject. A band pass filter (0.008–0.09 Hz) was used to reduce low-frequency drift and noise effects from the BOLD data.

### Functional connectivity analysis

Our data was first run through a seed-to-voxel bivariate correlation analysis, with PCC as the seed and FAs as a repeated measures variable. This was performed by using the CONN toolbox. An extraction of the BOLD time course from the seed (PCC and mPFC) and corresponding correlation coefficients of the BOLD time series for each voxel in the whole brain was carried out.

Voxel to voxel connectivity (r) versus voxel to voxel distance (mm) plot was used to evaluate the distribution of connectivity values (r). Global signal after denoising was regressed from the voxel BOLD timeseries. Distribution of connectivity values (r) significantly reduced after denoising. (See SI-3 for Quality assurance plots). Seed to voxel connectivity (z-transform normalized) differences between the different FA resting state sessions was investigated using two-sided paired t-tests within the CONN toolbox. These were done after confirming the main effect of flip angle using an ANOVA. For seed to voxel analysis using PCC as seed there were significant differences in precuneus. For the analysis using mPFC as a seed, there were significant differences in PCC, precuneus and posterior cingulate gyrus (See supplementary material SI-4). To account for multiple comparisons, we corrected p-value thresholds for a significant cluster (*p-corrected = 0*.*01*).

We identified the connectivity index values obtained in each FA block and compared the values between them. We checked for the difference in the functional connectivity between FA blocks using a paired samples t-test, Height threshold *p* < 0.001, cluster threshold *p* < 0.01, *p*-uncorrected and cluster size *p*-FWE corrected (min. 50 voxel cluster) was used.

We calculated FC using two seeds in separate seed to voxel analysis. These were picked apriori to be PCC and mPFC, this choice was based on their use in previous literature (Whitfield-Gabrieli & Ford, 2012). Given we interested in investigating the role of PCC in the default mode network under different FA scans, under the rationale that PCC is prone to physiological noise, a separate seed to voxel analysis with mPFC was used to cross-verify our functional connectivity results.

## Results

Before we made comparisons between different FA blocks or compared BOLD signals within and between blocks, we wanted to ensure that our dataset shows similar DMN activity as reported in previous literature when using the standard FA. To replicate standard DMN results with typically used parameters, we chose the 90° FA condition. We did a ROI-ROI based analysis to show that regions of PCC, mPFC and LP were functionally connected to each other. Moreover, the strength of this network was highest with PCC as the seed (See SI 1).

### Functional Connectivity results with PCC as seed

For the difference between functional connectivity in FA90° and FA15°, our results showed 5 clusters with significantly different connectivity strengths (See Fig 1(a)). Similarly, for the difference in functional connectivity between FA90° and FA50° there were 2 clusters that showed significantly different connectivity indices (See Fig 3(b)). No differences in functional connectivity were present between FA90° and FA77° blocks (See Fig 3(c)). Similarly, no differences in functional connectivity were found between FA50° and FA77° also (See Fig 3(e)). A comparison of FA77 and FA 15, showed one significantly different cluster (See Fig 3(f)). The details of the clusters are given in Table 1.

**Table 1:** The table lists the clusters with significantly different functional connectivity values between resting blocks using a seed to voxel analysis. No differences are reported for 90 vs. 77 and 77 vs. 50 blocks, as there were no significant differences found. The clusters thresholds were corrected for family wiser error with a threshold of 0.01 and only thus clusters with minimum 50 voxels have been reported.

| Cluster MNI coordinates (x, y, z) | T value | Maximum covered Voxel Area | Number of Voxels |
| --- | --- | --- | --- |
| -12 -46 +44 | 10.16 | Posterior cingulate gyrus/precuneus | 1302 |
| +12 -52 +26 | 6.77 | Posterior cingulate cortex | 128 |
| +44 -44 +16 | 5.88 | Posterior superior temporal gyrus | 89 |
| +24 +44 +36 | 9.21 | Medial prefrontal cortex | 69 |
| -34 -64 +50 | 5.39 | Precuneus | 115 |

PCC: 90-50

| Cluster MNI coordinates (x, y, z) | T value | Maximum covered Voxel Area | Number of Voxels |
| --- | --- | --- | --- |
| -48 -66 -20 | 6.44 | Lateral occipital cortex | 56 |
| +34 -74 +20 | 7.02 | Angular gyrus | 58 |

PCC: 50-15

| Cluster MNI coordinates (x, y, z) | T value | Maximum covered Voxel Area | Number of Voxels |
| --- | --- | --- | --- |
| -08 -62 +28 | 11.24 | Posterior cingulate gyrus | 532 |

PCC: 77-15

| Cluster MNI coordinates (x, y, z) | T value | Maximum covered Voxel Area | Number of Voxels |
| --- | --- | --- | --- |
| -06 -28 +30 | 8.38 | Posterior cingulate cortex | 1056 |

**Fig. 1.**
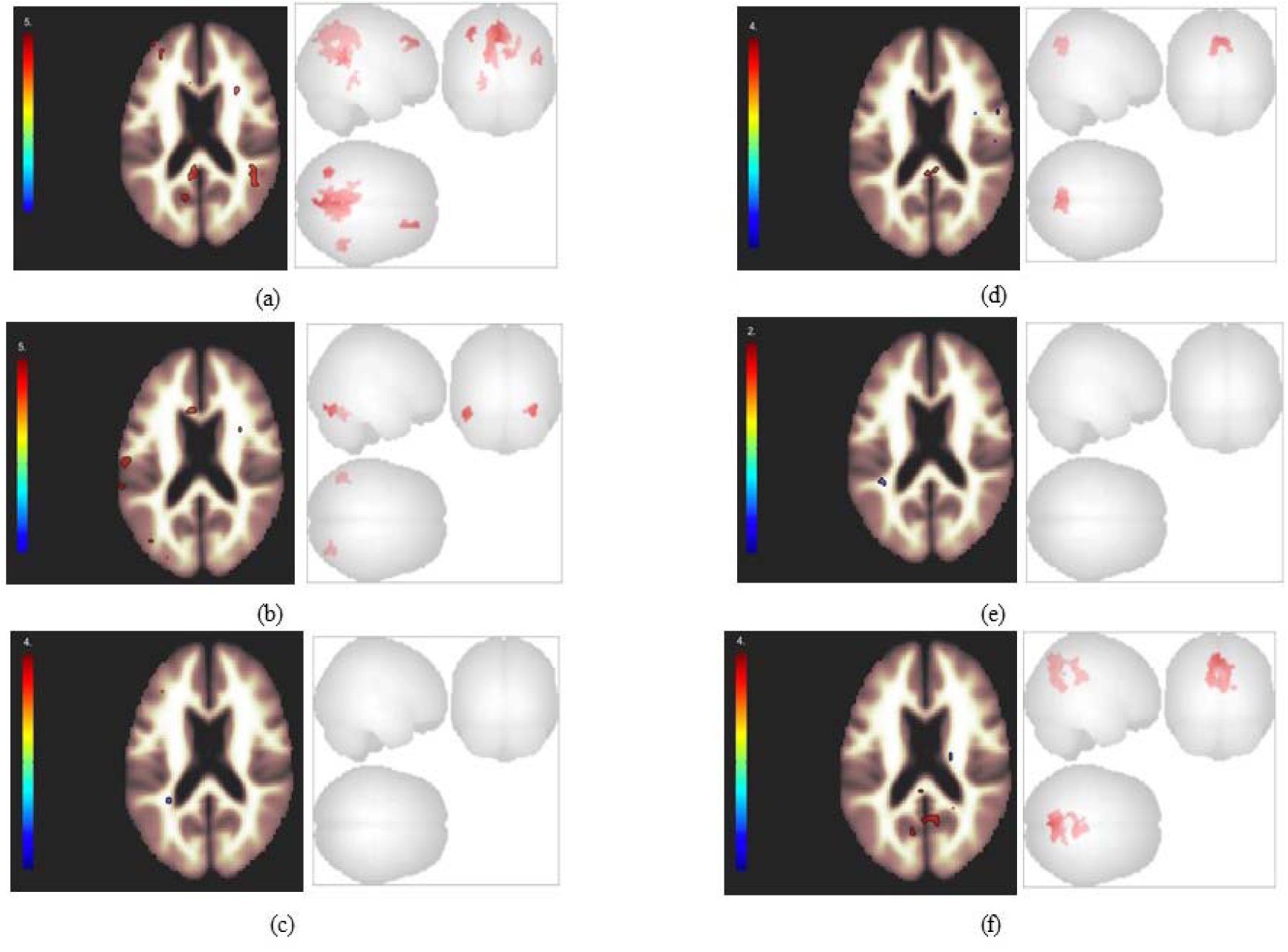
Functional connectivity difference between different flip angles after a seed to voxel analysis with PCC as a seed: (a) FA90° and FA15° (b) FA90° and FA50° (c) FA90° and FA77° (d) FA50° and FA15° (e) FA50° and FA77° and (f) FA77° and FA15° (Height threshold *p* < 0.001, cluster threshold *p* < 0.01 (cluster size p-FWE corrected).

### Seed to voxel connectivity, mPFC as seed

We reran the same bivariate seed-to-voxel correlation analysis for functional connectivity, but this time with medial prefrontal cortex (mPFC) as the seed. Since our hypothesised area of interest was the PCC, we wanted to explore differences in connectivity strength in this region independent of it being a seed. The same corrections and pre-processing steps were used as for with PCC as the seed. We also used the same corrections for multiple comparisons as in the previous analysis.

Our results again did not show any differences between FA77and FA50. However, we now found one significant cluster different between FA90 and FA77. There were 9 significantly different clusters in comparing connectivity indices for FA15 and FA90 blocks. Similarly, there were 10 significant different clusters identified in comparing FA77 and FA15 resting blocks. One significant cluster was identified as different between FA90 and FA50, and two clusters between FA50 and FA15 (See Table 2 and Figure 2).

**Table 2:**
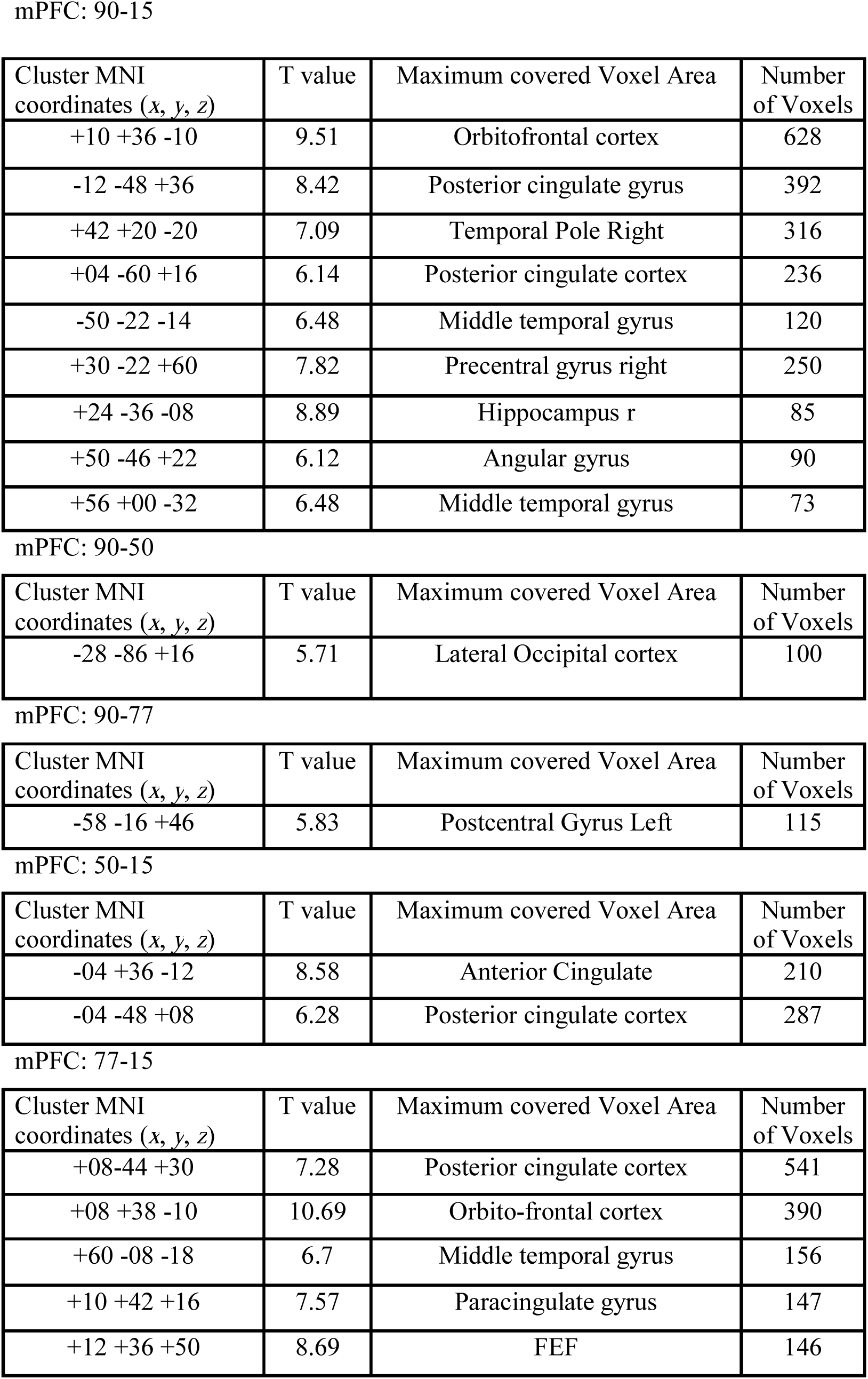
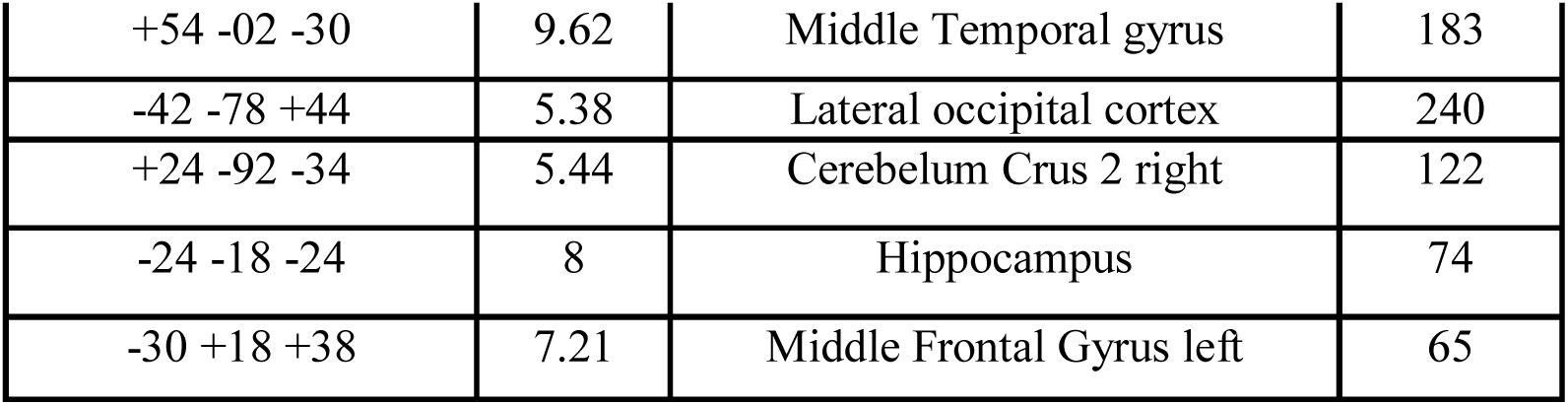
The table lists clusters found significantly different between resting blocks of different flip angles. The table does not list 77 vs. 50 resting blocks comparison, as there were no significant differences found in this comparison. The clusters reported here are after correction for multiple comparisons (Height threshold p < 0.001, cluster threshold p <0.01; minimum size of cluster was 50 voxels).

| Cluster MNI coordinates (x, y, z) | T value | Maximum covered Voxel Area | Number of Voxels |
| --- | --- | --- | --- |
| +10 +36 -10 | 9.51 | Orbitofrontal cortex | 628 |
| -12 -48 +36 | 8.42 | Posterior cingulate gyrus | 392 |
| +42 +20 -20 | 7.09 | Temporal Pole Right | 316 |
| +04 -60 +16 | 6.14 | Posterior cingulate cortex | 236 |
| -50 -22 -14 | 6.48 | Middle temporal gyrus | 120 |
| +30 -22 +60 | 7.82 | Precentral gyrus right | 250 |
| +24 -36 -08 | 8.89 | Hippocampus r | 85 |
| +50 -46 +22 | 6.12 | Angular gyrus | 90 |
| +56 +00 -32 | 6.48 | Middle temporal gyrus | 73 |

mPFC: 90-50
| Cluster MNI coordinates (x, y, z) | T value | Maximum covered Voxel Area | Number of Voxels |
| --- | --- | --- | --- |
| -28 -86 +16 | 5.71 | Lateral Occipital cortex | 100 |

mPFC: 90-77
| Cluster MNI coordinates (x, y, z) | T value | Maximum covered Voxel Area | Number of Voxels |
| --- | --- | --- | --- |
| -58 -16 +46 | 5.83 | Postcentral Gyrus Left | 115 |

mPFC: 50-15
| Cluster MNI coordinates (x, y, z) | T value | Maximum covered Voxel Area | Number of Voxels |
| --- | --- | --- | --- |
| -04 +36 -12 | 8.58 | Anterior Cingulate | 210 |
| -04 -48 +08 | 6.28 | Posterior cingulate cortex | 287 |

mPFC: 77-15
| Cluster MNI coordinates (x, y, z) | T value | Maximum covered Voxel Area | Number of Voxels |
| --- | --- | --- | --- |
| +08-44 +30 | 7.28 | Posterior cingulate cortex | 541 |
| +08 +38 -10 | 10.69 | Orbito-frontal cortex | 390 |
| +60 -08 -18 | 6.7 | Middle temporal gyrus | 156 |
| +10 +42 +16 | 7.57 | Paracingulate gyrus | 147 |
| +12 +36 +50 | 8.69 | FEF | 146 |
| +54 -02 -30 | 9.62 | Middle Temporal gyrus | 183 |
| -42 -78 +44 | 5.38 | Lateral occipital cortex | 240 |
| +24 -92 -34 | 5.44 | Cerebellum Crus 2 right | 122 |
| -24 -18 -24 | 8 | Hippocampus | 74 |
| -30 +18 +38 | 7.21 | Middle Frontal Gyrus left | 65 |

**Fig. 2.**
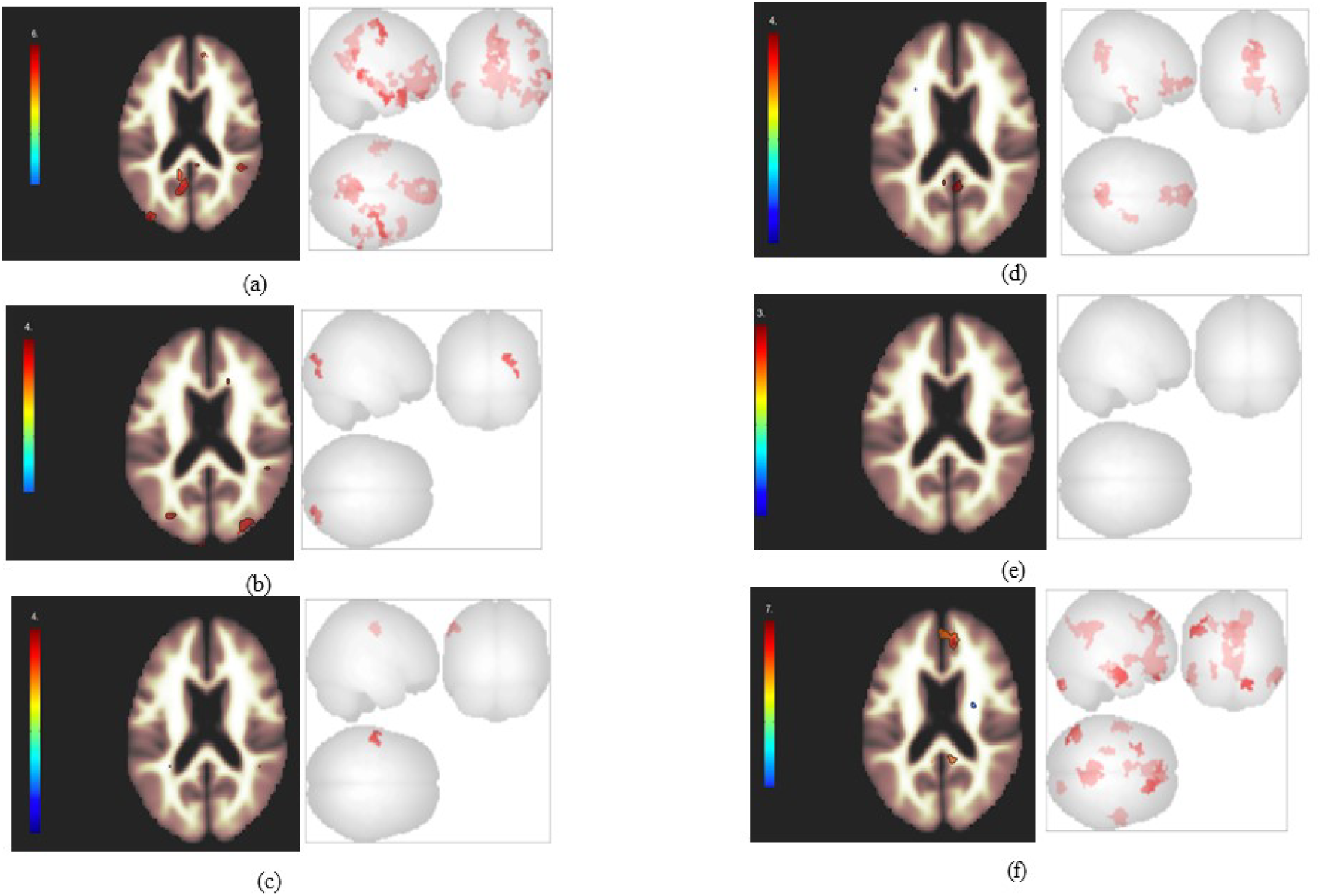
Functional connectivity difference between different flip angles after a seed to voxel analysis with mPFC as a seed: (a) FA90° and FA15° (b) FA90° and FA50° (c) FA90° and FA77° (d) FA50° and FA15° (e) FA50° and FA77° and(f) FA77° and FA15° (Height threshold *p* < 0.001, cluster threshold *p* < 0.01 (cluster size p-FWE corrected to minimum 50 clusters).

## Discussion

Our results raise concerns about fMRI activity recorded in PCC and precuneus during resting state. It has been shown that BOLD activity in these two areas do not show up across FAs (Renvall et al., 2014), when participants view a flickering checkerboard. We further show inconsistencies in BOLD signals recorded from these areas using a resting state fMRI recording. Similar to previous reports, these two areas succumb to changes in FAs.

Our findings provide an empirical criticism of the DMN. We show that the strength of the correlation between BOLD signals recorded while participants lay inside the scanner at rest, depends on the choice of FAs. Suspicions of physiological noise being present in the DMN have a long-standing history, for the network as whole (Shmueli et al., 2007; Munck et al., 2008; Murphy, Birn & Bandettini, 2013) and specifically with PCC as a node (Wise et al.,2004; Birn et al., 2006; Bianciardi et al., 2011). We provide empirical support to these previous findings by recording resting state activity using different FAs in counterbalanced blocks. This is important because lower FAs have been modelled and shown to reduce physiological noise, when thermal noise is controlled (Gonzalez-Castillo et al., 2011).

Our results show lots of clusters for which we had no predictions, i.e. areas outside the “core” default mode network. Our predictions were only for PCC and precuneus regions, since they could be independently verified in a flickering checkerboard task (Renvall et al., 2014) We do not attempt to interpret them here. But several of these areas like anterior cingulate cortex, hippocampus angular gyrus and temporal pole are also often reported as part of the DMN (Shulman et al., 1997; Buckner et al., 2008; Andrews-Hannah et al., 2010b). They could be independently investigated for physiological noise confounds with small flip angles in specific tasks that are reported to involve these regions. This is necessary before interpreting results for these regions from our study.

Our interpretations here are entirely driven from predicted differences in PCC and precuneus based on previous studies (Renvall et al., 2014). These regions are also prevalent in our data consistently in differences between high and low flip angle blocks, across seed to voxel analysis with two seeds. That is the differences in connectivity strength in these regions (PCC, precuneus and posterior cingulate gyrus) are present with both PCC and mPFC as a seed. Even though there are other clusters that are significantly different in comparing functional connectivity values between FA blocks, our interpretation here is hypothesis driven leaving us short of interpreting unexpected and unpredicted differences.

It could be that lower FAs reduce the quality of BOLD signals recorded in our study, but we see no reason as to why this would selectively affect PCC and precuneus while sparing other nodes of the DMN and other regions of the brain. This is also not true in the signal quality we get in the four different FA blocks, where signal quality is similar across the four blocks (se SI 3). Moreover, in our seed to voxel analysis (seed as PCC) we report no differences in functional connectivity between FAs 77 and 90 and no meaningful difference with mPFC as seed (i.e. no hypothesised differences). This further strengthens our claim that the reported DMN over the past two decades has been strongly influenced by the choice of FA, since most studies have used FAs between 77 and 90 from the prescribed Ernst angle of 77° (Greicius et al., 2003; Greicius et al., 2009; Fox et al., 2005;Fransson & Marrelec, 2008; Utevsky, Smith & Huettel, 2014; See also the protocol guidance document of the Human Connectome Project (2017), p.39).

We would also like to point out that our primary argument is not that mPFC or lateral parietal (LP) regions of the brain are part of the DMN. Our results are not inconsistent with such an interpretation. Whether mPFC and LP regions (or for that matter any other region of the brain) are part of the DMN are still open to both empirical and conceptual investigations. We do show inconsistencies with PCC and precuneus raising questions about their role in a putative DMN. The reported importance and role of both PCC and precuneus in the DMN might be confounded by noise given that rsfMRI scans have usually been done using FAs above 77 degrees.

Our results consistently show differences in PCC and precuneus regions of the brain with different flip angles, while participants are at rest inside the scanner. We get this consistent difference with two choices of seeds (PCC and mPFC) for a seed to voxel connectivity analysis. The results reported here challenge the robustness of the default mode network and its use for between group differences in different populations (i.e. elderly and clinically diagnosed). The latter because these differences are interpreted without removing confounds of physiological noise through FA alterations. Moreover, our results also generate scepticism for the use of PCC as the standard choice of seed for extracting the resting state time series, since the connectivity index of this region itself is prone to alterations with the flip angle.

Seen together with earlier studies (Renvall et al., 2014), there is a greater concern for previously reported BOLD activations in PCC and precuneus regions of the brain both during rest and task. PCC has been deemed responsible for multiple functions including emotional processing (Maddock, Garrett & Buonocore, 2002), conscious experiences (Vogt & Laureys, 2006), and regulating and directing attention (Hahn, Ross & Stein, 2007). Precuneus has been linked with self-related processing, autobiographical memories and realization of conscious experiences (Cavanna, 2007). These findings in the light of our work and earlier studies (Renvall et al., 2014) are left vulnerable to the choice of FAs. Our result that the choice of FA alters the strength of PCC involvement is more critical because PCC activations constitute a widely reproducible result in studies on DMN.

It may be the case that the results of some studies truly reflect PCC and precuneus involvement; however, their claims are open to criticisms. We suggest, in line with Renvall et al (2014), that fMRI studies be done over a variety of FAs to produce reliable results, at least if one is interested in showing involvement of PCC and precuneus in task-related and rest-related designs. Here we present data to show that FA choices alter connectivity strengths in the default mode network, largely in the regions of the posterior cingulate cortex, posterior cingulate gyrus and precuneus. Our results seen through the lens of physiological noise minimization provide a novel empirical criticism of the findings on default mode network.

## Acknowledgements

The study reported here was conducted at the National Neuroimaging Facility at CBCS, University of Allahabad. We thank the Department of Science & Technology (DST), Government of India for the grant (SR/CSRI/01/2015) to set up the facility and make this research possible.

## Author Contributions

IS and NS designed and conceptualised the study. IS and AK collected the data. AK and IS analysed the data. IS, NS, and AK interpreted the results and wrote the manuscript.

