## Supplementary Material for "Default(y) Mode Network: Important regions of DMN do not survive alterations in flip angles"

**Supplementary Information**

**SI 1** Connectome Ring analysis for the standard DMN using 90° FA.

We replicate the existing DMN using ROI to ROI analysis, and show using PCC as a seed produces strongest results in functional connectivity in the network.

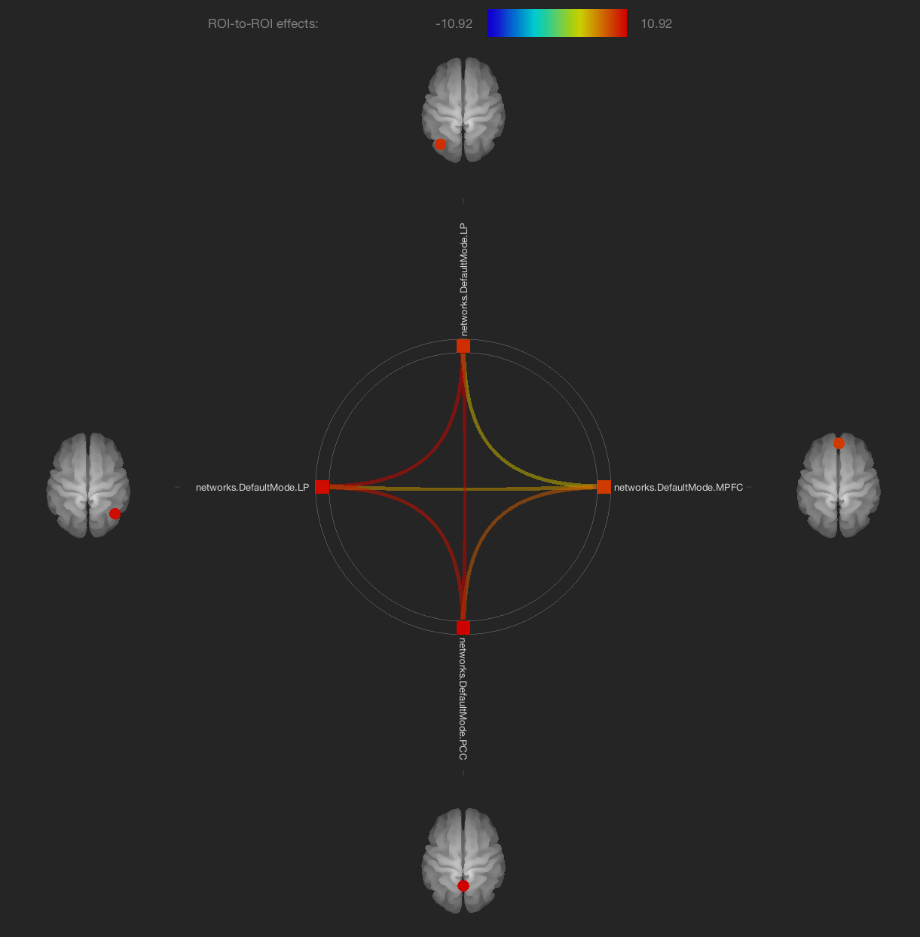

**Fig S1.** Connectome ring: ROI to ROI effects for DMN analysis. The four ROIs (nodes of the DMN) are PCC, mPFC and left and right LP areas. Our results indicate strong functional connectivity between the four regions.

**Tables S1.** DMN statistic showing connectivity difference (the difference is reasonably significant for PCC). The results show strong functional connectivity in the DMN, and the connectivity is the strongest when PCC is used as a seed in multiple ROI-ROI analyses. This replicates exisiting findings in the DMN at 90° FA.

Results when using Lateral Parietal (L) region to calculate functional connectivity within the pre-defined default mode network. *F*(3,13) = 89.85, *p* <0.001

| Connection analysed | T-value | *p*-FDR | *p*-FWE |
| --- | --- | --- | --- |
| Lateral Parietal (L) - PCC | 10.92 | <0.001 | <0.001 |
| Lateral Parietal (L) – Lateral Parietal (R) | 10.64 | <0.001 | <0.001 |
| Lateral Parietal (L) - mPFC | 4.44 | <0.001 | <0.001 |

Results when using PCC region to calculate functional connectivity within the pre-defined default mode network. *F*(3,13) = 76.08, *p* <0.001

| Seed-to-ROI | T-value | *p*-FDR | *p*-FWE |
| --- | --- | --- | --- |
| PCC- Lateral Parietal (L) | 10.92 | <0.001 | <0.001 |
| PCC- Lateral Parietal (R) | 10.36 | <0.001 | <0.001 |
| PCC- mPFC | 7.75 | <0.001 | <0.001 |

Results when using Lateral Parietal (R) region to calculate functional connectivity within the pre-defined default mode network. *F*(3,13) = 59.69, *p* <0.001

| Seed-to-ROI | T-value | *p*-FDR | *p*-FWE |
| --- | --- | --- | --- |
| Lateral Parietal (R) - PCC | 10.36 | <0.001 | <0.001 |
| Lateral Parietal (R) – Lateral Parietal (L) | 10.64 | <0.001 | <0.001 |
| Lateral Parietal (R) - mPFC | 5.74 | <0.001 | <0.001 |

Results when using mPFC region to calculate functional connectivity within the pre-defined default mode network. *F*(3,13) = 37.21, *p* <0.001

| Seed-to-ROI | T-value | *p*-FDR | *p*-FWE |
| --- | --- | --- | --- |
| mPFC-PCC | 7.75 | <0.001 | <0.001 |
| mPFC- Lateral Parietal (R) | 5.74 | <0.001 | <0.001 |
| mPFC- Lateral Parietal (L) | 4.44 | <0.001 | <0.001 |

**SI 2** The figure shows a 2-D slice-wise section of the BOLD activity under four different FA resting blocks, when calculated using a ROI to ROI analysis. Panel (a) shows it for FA 15, panel (b) is for FA 50, (c) and (d) are for FA77 and FA 90 respectively.

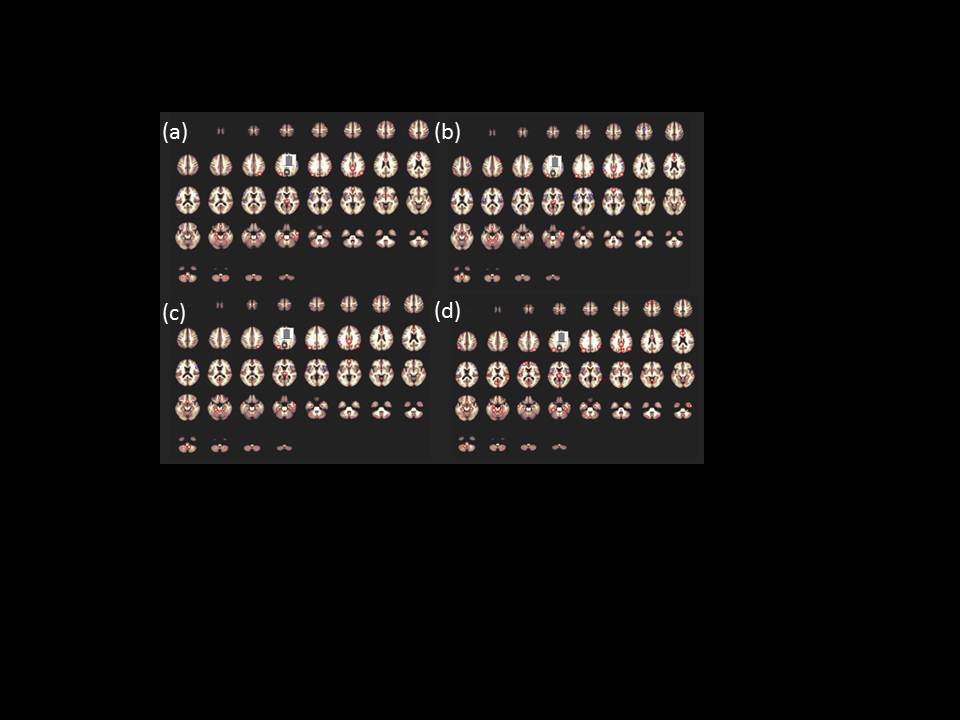

**SI 3:** Given below are quality assurance plots for our recorded BOLD activity in the four different FA blocks. There is reasonably similar quality in all four sessions, addressing any concerns for signal-to-noise ratio being the confound in our results.

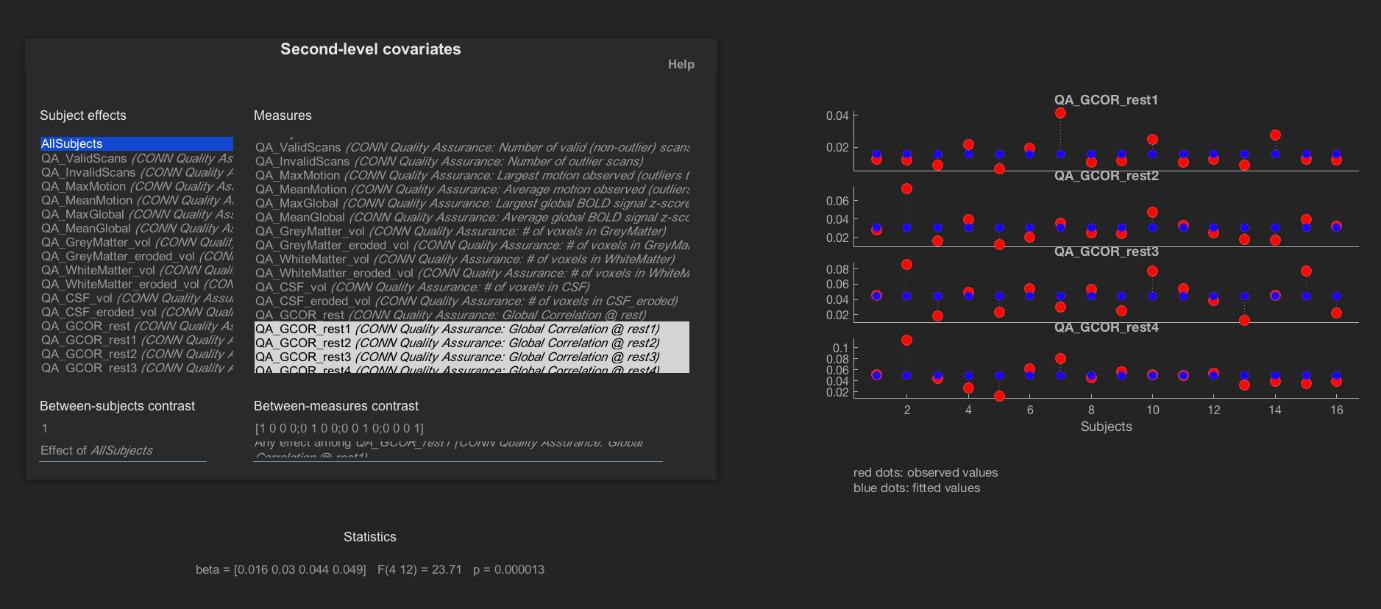

**Fig**. Quality assurance (QA) plot for global correlation between four different FA values for all subjects (rest 1 for FA15^o^, rest 2 for FA50^o^, rest 3 for FA77^o^ and rest 4 for FA90^o^). The corresponding beta values are 0.016, 0.030, 0.040 and 0.049 respectively (F-statistics, p=0.000013).

S4: One-way ANOVA results with flip angle as the independent variable

Table S4.1: Clusters identified in a one-way ANOVA (PCC seed) at p<0.01 cluster threshold. The results are corrected using p-FWE and only those clusters that had more than 50 voxels are reported.

| Cluster MNI coordinates (*x*, *y*, *z*) | Maximum covered Voxel Area | Number of Voxel |
| --- | --- | --- |
| -02 -49 +61 | Precuneus | 528 |
| +26 +30 +42 | Middle Frontal Gyrus | 112 |

Table S4.2: Clusters identified in a one-way ANOVA (mPFC seed) at p<0.01 cluster threshold. The results are corrected using p-FWE and only those clusters that had more than 50 voxels are reported.

| Cluster MNI coordinates (*x*, *y*, *z*) | Maximum covered Voxel Area | Number of Voxel |
| --- | --- | --- |
| -2 -44 +38 | Precuneus | 537 |
| +52 -62 +28 | Angular Gyrus | 91 |
| +11 -42 +26 | Posterior Cingulate Cortex | 243 |
| 11 -35 +42 | Posterior Cingulate Gyrus | 528 |
